# Noninvasive Ultrasonic Glymphatic Induction Enhances Intrathecal Drug Delivery

**DOI:** 10.1101/2020.10.21.348078

**Authors:** Muna Aryal, Quan Zhou, Eben L. Rosenthal, Raag D. Airan

**Author notes:** Corresponding Author and Lead Contact: Raag D. Airan MD, PhD.

## Abstract

Intrathecal drug delivery is routinely used to bypass the blood-brain barrier in treating varied central nervous system conditions. However, the utility of intrathecal delivery is limited by poor parenchymal uptake of agents from the cerebrospinal fluid. We demonstrate that a simple noninvasive transcranial ultrasound protocol significantly increases the brain parenchymal uptake of intrathecally administered drugs and antibodies. Essentially, we show that our protocol of transcranial ultrasound can accelerate glymphatic fluid transport from the cisternal space into the parenchymal compartment. Specifically, we administered small (∼1kDa) and large (∼150 kDa) molecule agents into the cisterna magna of rats and then applied low, diagnostic-intensity focused ultrasound in a scanning protocol throughout the brain. Using both real-time magnetic resonance imaging and ex vivo histologic analyses, we observed significantly increased uptake of each agent into the brain parenchyma from the cisternal cerebrospinal fluid, notably with no brain parenchymal damage. The low intensity of the ultrasound and its noninvasiveness underscores the ready path to clinical translation of this technique for whole-brain delivery of a variety of agents. Furthermore, this technique can be used as a means to probe the causal role of the glymphatic system in the variety of disease and physiologic processes to which it has been correlated.

**eTOC Summary:** A translation-ready ultrasound technique enhances the brain penetration of intrathecally delivered agents via upregulating the glymphatic pathway.

## Introduction

Drug delivery to the brain is significantly limited by the blood-brain barrier (BBB), which excludes ∼98% of potential small molecule therapeutics and nearly 100% of large therapeutics (Abbott and Romero, 1996; Neuwelt et al., 2008; Pardridge, 2005). In principle, if an agent is administered into the cerebrospinal fluid (CSF) of the cisterns or ventricles of the central nervous system (CNS), e.g. via intrathecal delivery during a spinal tap, the agent would already be across the BBB and therefore able to access the brain and spine parenchyma. Indeed, intrathecal delivery is already used in the treatment or prophylaxis of a variety of CSF-based diseases, including leptomeningeal metastatic cancer and infectious meningitis (Jain et al., 2019). However, drug penetration into the brain and spine parenchyma from the CSF is known to be severely limited (Burch et al., 1988; Calias et al., 2014; Leal et al., 2011). A means to overcoming this effective CSF-parenchyma barrier could greatly expand the utility of myriad off-the-shelf therapeutics for the treatment of numerous CNS diseases.

Recently, researchers have observed that vascular pulsations may drive active transport of cisternal CSF fluid into the interstitial compartment of the brain parenchyma, a system coined the “glymphatic pathway” (Iliff et al., 2012, 2013). While the glymphatic pathway could be utilized for drug delivery, at baseline its rate of fluid transport is insufficient to drive significant convection of intrathecally administered agents into the brain parenchyma (Rasmussen et al., 2018). Further, while the glymphatic system has been linked to a variety of physiological states, like sleep, and diseases like Alzheimer’s disease or traumatic brain injury, these studies are fundamentally correlative as there are no described means for independently controlling glymphatic transport (Ahn et al., 2019; Benveniste et al., 2019; Gaberel et al., 2014; Goulay et al., 2017; Hawkes et al., 2014; Iliff et al., 2012, 2014; Jessen et al., 2015; Kress et al., 2014; Louveau et al., 2015; Mesquita et al., 2018; Peng et al., 2016; Rasmussen et al., 2018; Yang et al., 2013).

As the glymphatic system is driven by convective pressures induced by arterial pulsation (Iliff et al., 2013), and since ultrasound is a high-frequency wave of pressure oscillations in the medium, we hypothesized that ultrasound application could induce similar pressure oscillations in the perivascular and interstitial space as seen with glymphatic transport, and that this could be used to increase the brain parenchymal penetration of intrathecally administered agents. Several groups have shown that ultrasound may increase the diffusion of agents within tissue after either pairing ultrasound with exogenous microbubble contrast agents (Aryal et al., 2015; B et al., 2019; Chen and Konofagou, 2014; Chen et al., 2014; Etame et al., 2012; Wei et al., 2013) or by using high *in situ* pressures (Curley et al., 2020; Mead et al., 2019; Mohammadabadi et al., 2020). It has yet been determined whether a low-intensity ultrasound protocol on its own may increase the cisternal CSF-parenchymal transport that is the hallmark of the glymphatic pathway (Iliff et al., 2013). Here, we demonstrate that we may indeed use noninvasive transcranial low-intensity ultrasound to increase the parenchymal penetration of intrathecally administered small and large therapeutic agents via glymphatic upregulation.

## Results

Given the known effects of anesthesia and sleep on glymphatic transport (Gakuba et al., 2018; Mendelsohn and Larrick, 2013), titration of isoflurane anesthetic dose and environmental heating was used with cardiorespiratory monitoring to ensure physiologic stability during the up to 4 hours of each experiment and intervention, with respiratory rates maintained in the range of 50 to 60 breaths per minute, heart rates of approximately 300–370 beats per minute, O_2_ saturation of approximately 98%–100%, and body temperature of 36.5°C to 37.5°C. A scanning ultrasound protocol was chosen to treat the whole rat brain with transcranial focused ultrasound of an intensity below FDA-approved limits for diagnostic ultrasound (0.25 mechanical index, MI, *in situ*, 7.7 % local duty cycle, for a total of 10 min.; **Fig. 1**). This intensity of ultrasound was chosen as it is less than or similar to the intensities used in routine diagnostic ultrasound imaging of adult and neonatal human patient brains, and is readily achievable with ultrasound systems designed for diagnostic or therapeutic transcranial ultrasound applications in the adult human brain (Mainprize et al., 2019). Notably, the total temperature rise in the sonicated zone due to this level of ultrasound exposures is estimated to be <0.01 °C (Nyborg 1988; Haar and Coussios 2007).

**Fig. 1.**
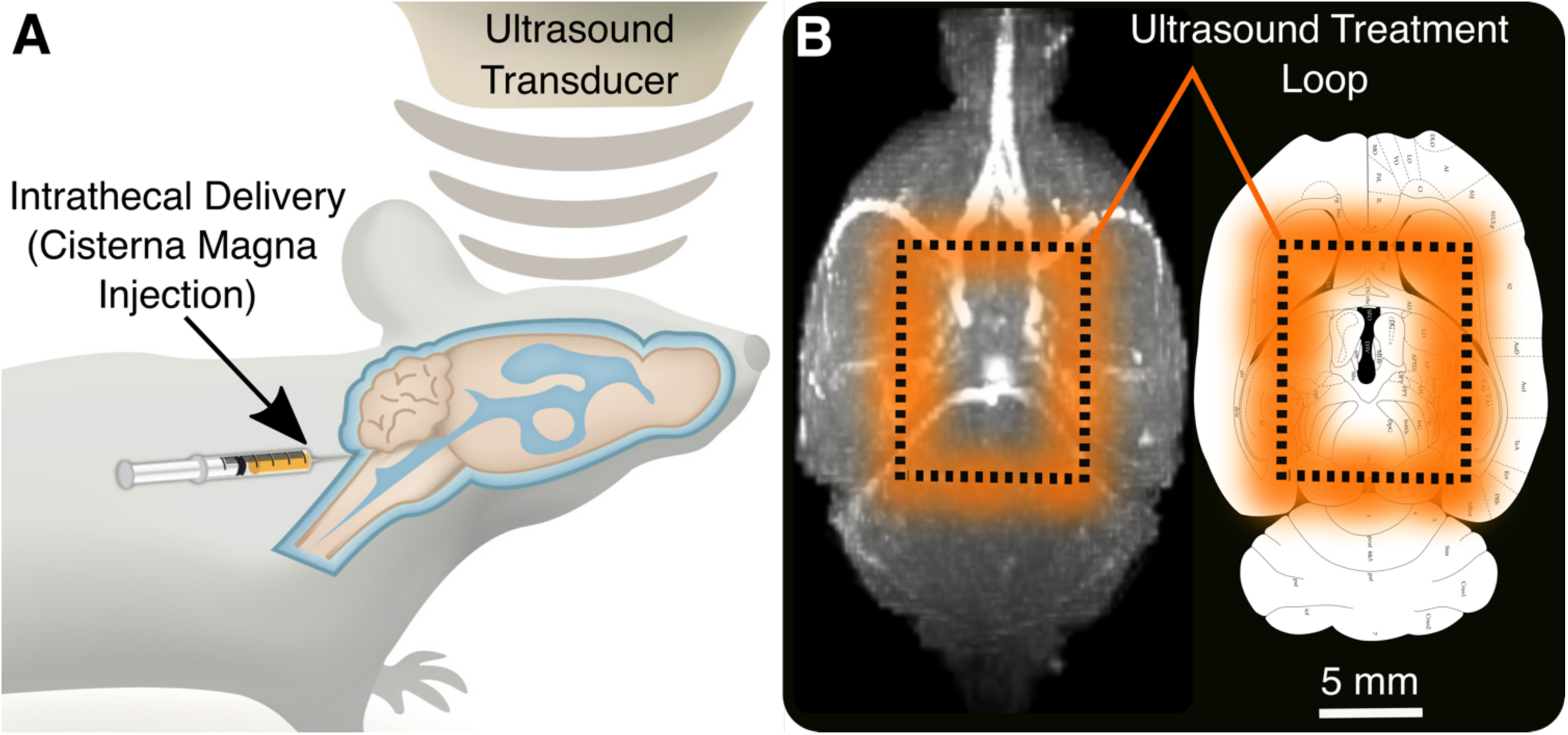
Schematic of ultrasonic glymphatic induction for enhancing the brain penetration of an intrathecally administered agent. **A**. Following intrathecal injection of an agent into the cisternal cerebrospinal fluid (CSF) to bypass the blood-brain barrier (BBB) transcranial focused ultrasound (FUS) is applied across the intact skull. **B**. Scanning an ultrasound focus across the whole brain is hypothesized to increase glymphatic transport of cisternal CSF into the brain. The ultrasound focus trajectory is indicated by the dashed line, with the full-width at half maximum of the ultrasound field in orange, and is overlaid onto a maximum intensity projection (MIP) of a 3D volumetric T1w MRI (*left*; major cerebral arteries in white) and a representative transverse Paxinos brain atlas section (*right*) (Paxinos 2017), indicating the relevant anatomical structures. Ultrasound protocol: 650 kHz ultrasound frequency, 0.25 MI (0.2 MPa estimated *in situ* peak negative pressure), held continuously on while scanning the indicated 8 × 10 mm rectangular trajectory repeatedly for 10 min; estimated local duty cycle of 7.7% given the focus transverse full-width at half-maximum (FWHM) of 2.78 mm (longitudinal FWHM 12 mm); 24 sec per 36 mm loop, for 25 loops across 10 min.

### Transcranial ultrasound noninvasively accelerates glymphatic transport of a 1 kDa tracer

A gadolinium (Gd)-chelate MRI contrast agent was injected intrathecally into the cisterna magna, to label the CSF to allow MRI visualization of glymphatic CSF transport from the basal cisterns into the brain parenchyma. 3D T1w MRI images revealed that perivascular influx of CSF into the brain is observed initially 12 min after intrathecal Gd-chelate administration. Dynamic quantitative T1-mapping MRI imaged the time-dependent CSF influx, parenchymal uptake, and clearance of the Gd-chelate (**Fig. 2B**). Without ultrasound intervention, the 1 kDa Gd-chelate enters the brain from the cisterns with a peak brain concentration at approximately 35 min and then clears from the brain interstitial compartment within 3 h from injection (**Fig. 2B**). With a low-intensity transcranial scanning ultrasound treatment, a more diffuse pattern of Gd-chelate brain distribution was observed. The peak parenchymal uptake with ultrasound was increased by 72-101% with a relatively delayed peak of 105 min from injection, and with increased tracer in the brain at 3 h post-injection (**Fig. 2; Supplementary Video V1**). Notably, the contrast agent entered the brain preferentially near sites of arterial influx into the parenchyma (**Fig 1B**, *left*; **Fig. 2**), in keeping with known patterns of glymphatic entry into the brain (Iliff et al., 2013). We noted a statistically significant difference of brain parenchymal tracer uptake in these trends between the ultrasound and sham conditions at 35, 70, and 105 min following tracer administration.

**Fig. 2.**
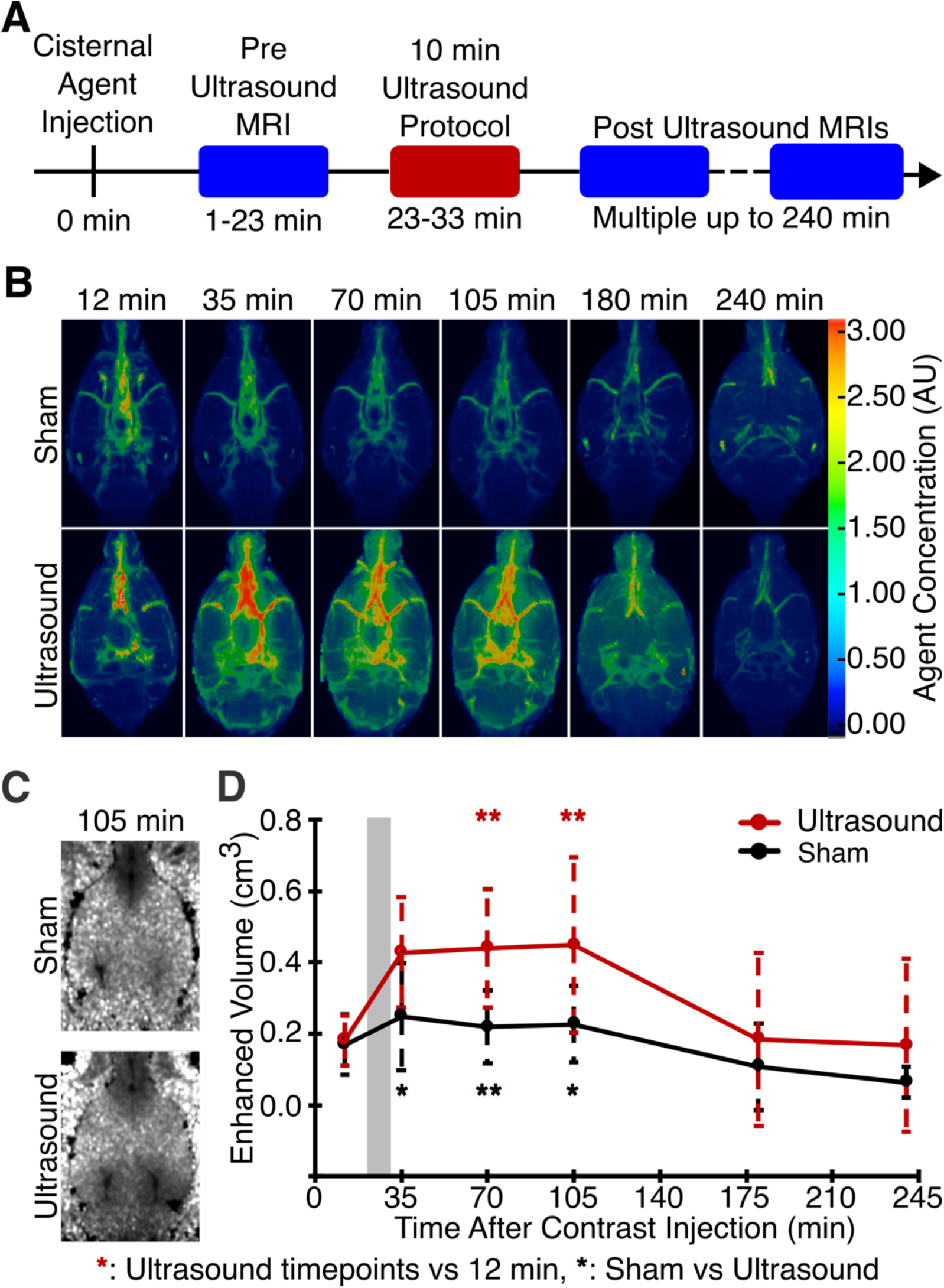
Transcranial ultrasound noninvasively accelerates glymphatic transport into the brain parenchyma for an intrathecally administered 1 kDa MRI tracer. **A**. Experimental timeline to measure the spread of intrathecally administered MRI contrast agent into the brain using quantitative T1-mapping MRI before and after ultrasound intervention. **B**. Representative pseudocolor T1w MRI images of rat brains following intrathecal MRI contrast agent injection with sham (*top*) or ultrasound (*bottom*) intervention. **C-D**. Quantitative T1-mapping MRI was used to quantify the contrast agent concentration in the brain before and after intervention. **C**. Representative T1-maps showing contrast agent in the brain with (*bottom*) and without (*top*) transcranial ultrasound application, at 105 min after contrast agent administration and 70 min from the ultrasound intervention; dark regions indicate higher MRI contrast agent concentration. **D**. Brain volume containing a resolvable amount of the intrathecally administered contrast agent over time showing that the 1 kDa contrast agent is driven into a significantly larger volume of brain with the ultrasound intervention, versus sham. Gray column indicates ultrasound intervention timing. *Presented as mean ± S*.*D. for groups of n = 10-11 (12 -105 min) and n = 3-4 (180 - 240 min). *: p ≤ 0*.*05, **: p ≤ 0*.*01 by ANOVA and post-hoc t-tests, comparing ultrasound to sham (black asterisk) and comparing ultrasound timepoints with the baseline at 12 min (red asterisk). Only the significantly different comparisons are noted*.

### Ultrasound noninvasively increases glymphatic transport of both small and large agents

To gain higher resolution of the distribution of the delivered agent, during Gd-chelate injection, we co-administered an optical (infrared-fluorescent) dye (∼1 kDa, IRDye800CW) in free form or conjugated to the therapeutic antibody panitumumab (∼150 kDa, in active clinical use for EGFR-targeted therapy) (Gao et al., 2018) to model the delivery of both small and large therapeutic agents. Glymphatic upregulation with this ultrasound protocol was confirmed as before using MRI visualization of the Gd-chelate. Animals were sacrificed two hours after agent administration (at the peak parenchymal uptake noted in the initial experiments, **Fig. 2D**). Histologic infrared fluorescent microscopy verified that both the small and large optical tracers indeed penetrated to a greater degree into the brain with this brief, 10 min noninvasive ultrasound therapy (**Fig. 3**).

**Fig. 3.**
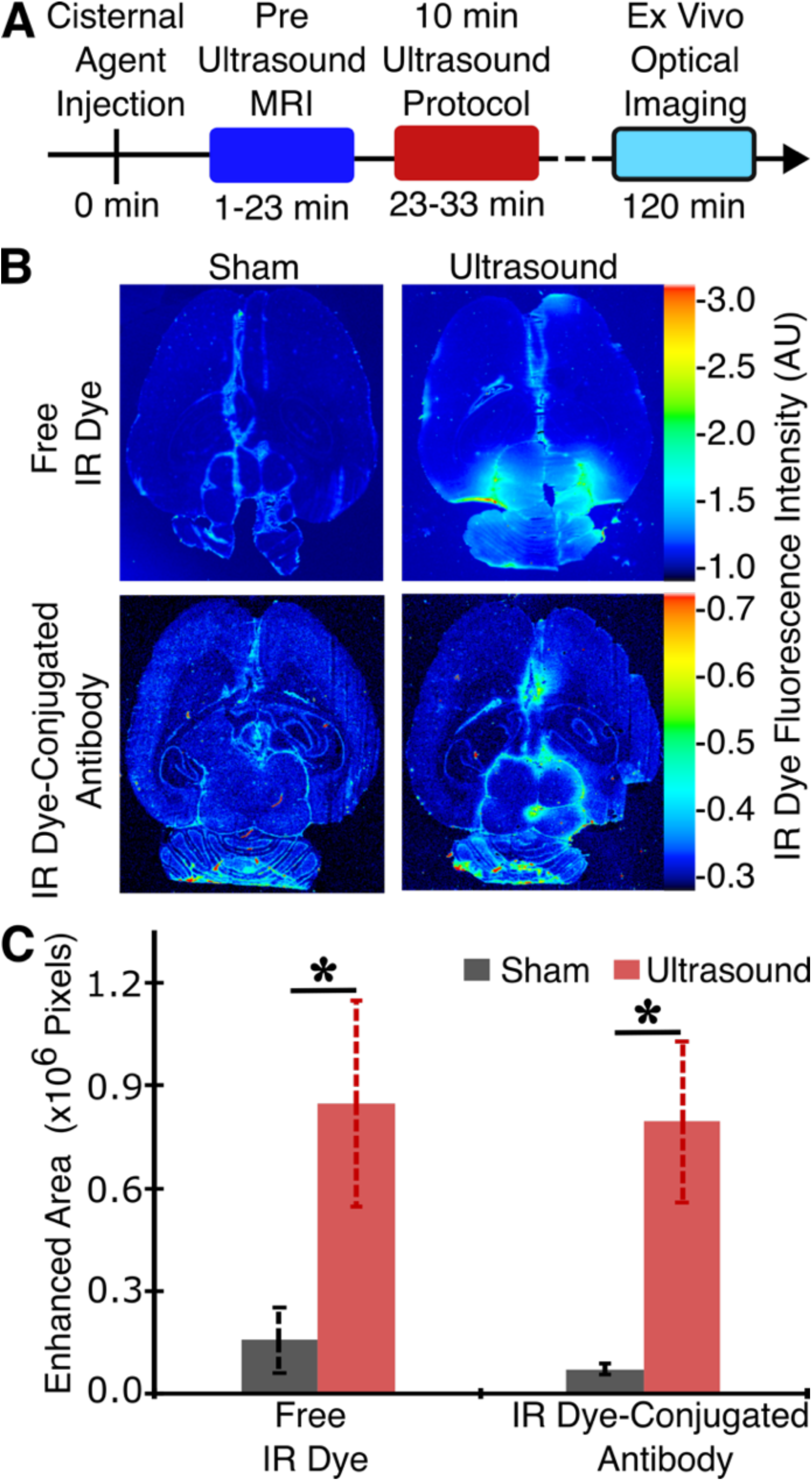
Transcranial ultrasound noninvasively accelerates glymphatic transport into the brain of small (1 kDa) and large (150 kDa) molecule agents following intrathecal administration. **A**. Experimental timeline to measure ultrasound-induced changes in brain penetration of intrathecally administered small and large optical agents. **B**. Representative pseudocolor near-infrared images of brain slices following cisternal injection of a small molecule optical tracer (*top*; ∼1 kDa, IRDye800CW dye) and a large molecular tracer (*bottom*; ∼150 kDa, Panitumumab-IRDye800). **C**. Near-infrared fluorescent imaging-defined dye-enhanced area for the small and large molecular tracers revealed that both agents penetrated into the brain to a significantly greater degree with ultrasound, compared to sham. *Presented as mean ± S*.*D. for groups of n = 3-4 for each agent. *: p ≤ 0*.*05 by two-tailed t-tests, comparing ultrasound to sham*.

### Ultrasonic glymphatic induction is safe

To evaluate the safety of this approach, high-field 7T MRI and histologic evaluation were utilized in both the acute and delayed settings, up to 72 h following ultrasound intervention. No evidence of microhemorrhage or edema was noted using T2*w MRI either within hours of intervention (**Fig. 4**) or up to 72 h following intervention (**Fig. S2**). Notably, no prolonged Gd-chelate deposition was seen in the brain up to 72 h, by when the CSF is known to be fully replaced (**Fig. S2**) (Lee et al., 2018; Liu et al., 2004). High-field histological evaluation (**Fig. 4C-E)** confirmed the lack of brain parenchymal damage with this intervention. Importantly, the total temperature change in the sonicated zone with this ultrasound protocol (0.25 MI *in situ*, 7.7% local duty cycle for 10 min) is estimated to be <0.01 °C based on the bio-heat transfer equation (Nyborg 1988; Haar and Coussios 2007). Further, this *in situ* intensity of ultrasound is similar to that used commonly for diagnostic brain imaging in both adult and neonatal human populations, and is well below FDA guidelines for ultrasound application in human tissue (Nelson et al., 2009). Therefore, this level of safety with this approach is expected.

**Fig. 4.**
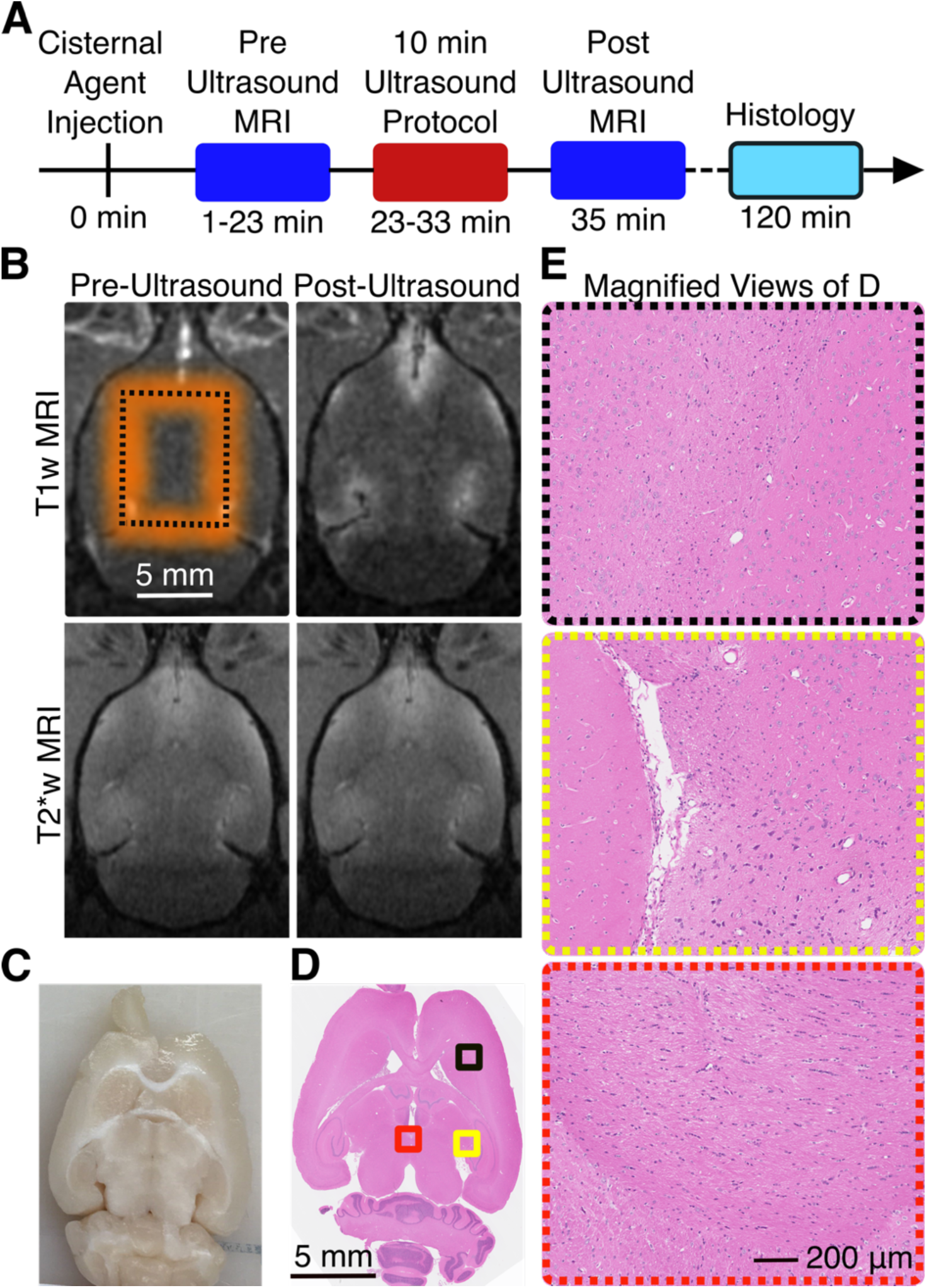
Ultrasonic glymphatic induction is safe. **A**.Experimental timeline for safety assessment. **B**. Representative MRI images showing no signs of damage, including edema or hemorrhage, in the brain parenchyma before (*left*) and after (*right*) transcranial ultrasound application (trajectory in orange; 0.25 MI in situ, 7.7% local duty cycle for 10 min). **C-E**. *Ex vivo* brain slice analysis. **C**. Representative bright-field image and **D**. Representative hematoxylin and eosin-stained (H & E) transverse sections of the brain of the same rat as in **B. E**. Magnified views of the indicated areas of **D** demonstrating no evidence of brain parenchymal damage with this ultrasound protocol.

## Discussion

We have demonstrated that low-intensity noninvasive transcranial ultrasound can upregulate the glymphatic pathway to improve the efficacy of intrathecal drug delivery. With MRI, we observed that this ultrasound protocol safely accelerates the transport of a ∼1 kDa MRI tracer from the CSF into the interstitial space before it clears from the brain **(Fig. 1, 2, S1, Video V1, Table S1)** with a clearance timeline of 3 hrs. Using optical tracers, we validated the MRI findings using a ∼1 kDa optical tracer that has a similar molecular weight as the MRI tracer and which models the distribution of small molecule drugs that are commonly intrathecally administered, like methotrexate (**Fig. 3**). Further, we used the same optical probe conjugated to a ∼150 kDa therapeutic antibody panitumumab (Gao et al., 2018) and saw similar increases of brain parenchymal uptake of this larger therapeutic agent (**Fig. 3**). Importantly, we saw no evidence of brain parenchymal damage with this approach (**Fig. 4, S2**).

Overall, our results suggest that low-intensity noninvasive transcranial ultrasound may be used to increase the whole-brain delivery of a variety of off-the-shelf small or large therapeutic agents, following the same intrathecal administration that is used routinely in clinics worldwide to administer therapeutic agents into the CSF (Burch et al., 1988; Calias et al., 2014; Edeklev et al., 2019; Eide et al., 2018; Jain et al., 2019; Leal et al., 2011; Liu et al., 2004; Mack et al., 2016). Further, this method provides a means to directly upregulate glymphatic transport, which could be used for causative evaluation of the role of the glymphatic system in the variety of physiological and disease processes to which the glymphatic system has been correlated (Abbott et al., 2018; Ahn et al., 2019; Aspelund et al., 2015; Benveniste et al., 2019; Gaberel et al., 2014; Gakuba et al., 2018; Iliff et al., 2014; Kress et al., 2014; Lundgaard et al., 2017; Mader and Brimberg, 2019; Mendelsohn and Larrick, 2013; Peng et al., 2016; Plog and Nedergaard, 2018; Pu et al., 2019; Simon and Iliff, 2016). Given the low intensity of ultrasound necessary for these results (Bein et al., 2006; Chen et al., 2018; Meijler and Steggerda, 2019; Naqvi et al., 2013; Nelson et al., 2009; Onweni et al., 2020; Purkayastha and Sorond, 2012; Rasulo et al., 2017; Sarkar et al., 2007), at levels readily achievable with currently-utilized clinical transcranial ultrasound systems (Abrahao et al., 2019; Arvanitis et al., 2013; Bystritsky et al., 2011; Fini and Tyler, 2017; Ghanouni et al., 2015; Hersh et al., 2016; Hynynen and McDannold, 2004; Izadifar et al., 2020; Jolesz, 2009; Lamsam et al., 2018; Medel et al., 2012; Poon et al., 2017; Pouliopoulos et al., 2020; Vykhodtseva et al., 2008), and the absence of non-therapeutic exogenous agents in this protocol or the need for chemical modification of the drug to be delivered, there is a ready path for clinical translation of a therapy based on these results.

## Methods and Materials

### Animals

The Institutional Animal Care and Use Committees of Stanford University approved all animal experiments. Tests were performed in 42 male Long-Evans rats with bodyweight 300–350 gm (Charles River Laboratories, Wilmington, MA, USA). Animals were randomly assigned to one of two groups: (1) no treatment (Sham), and (2) treatment (Ultrasound). Ultrasound (0.25 MI *in situ*, 7.7% duty cycle for 10 min) or sham was applied transcranially throughout the brain (**Fig. 1A**). Before each procedure, the fur on the neck was shaved and a cisterna magna injection of a gadolinium (Gd) chelate (Multihance, Bracco Diagnostics, NJ, USA) was performed while the animal was anesthetized under isoflurane. The body temperature, cardiac and respiratory rates, and O_2_ saturation were monitored throughout the experiment and the isoflurane level was titrated to keep these parameters constant; environmental heating was used to help maintain body temperature. Localizer, FLASH-T1-3D, T1-mapping, and T2*-weighted MR images were taken to visualize glymphatic transport across the brain, to quantify Gd-chelate kinetic parameters, and to evaluate for parenchymal damage. In separate cohorts, either of two different sized optical tracers, a small molecule (IR800CW Carboxylate, LICOR, Lincoln, NE, USA; ∼1 kDa) or a large molecule (Panitumumab-IRDye800: ∼150kDa, 5 nM, produced under GMP at the Leidos Biomedical Research Center, Frederick, MD, USA) (Gao et al., 2018) were co-delivered with the Gd-chelate to model the delivery of similar-sized therapeutic agents.

### Intrathecal Cisterna Magna Injection

For anesthesia, the animals were induced with 5% isoflurane in oxygen using an induction chamber and then switched to a maintenance dose of 2%. The animal was positioned in a stereotaxic frame (Stoelting, Wood Dale, IL, USA), immobilized with ear bars, and then the head flexed to 45 degrees (Liu and Duff, 2008; Santos et al., 2018). A 27-gauge catheter (Butterfly Needle, SAI Infusion Technology, Lake Villa, IL, USA) was inserted in the cisterna magna to inject up to 80 µl of tracers (Gd-chelate: MultiHance, gadobenate dimeglumine; Bracco Diagnostics Inc, NJ, USA; 0.21 ml/kg) slowly over 30 seconds. To model the delivery of similar-sized therapeutic agents, two different sized molecules, free dye (IRDye800CW Carboxylate, LICOR, Lincoln, NE, USA; ∼1 kDa; 36 nmol/kg) and IR dye-conjugated antibody (Panitumumab-IRDye800: ∼150 kDa; 0.133 mg/kg) were co-delivered with the Gd-chelate. The respiratory rate (45-50 breaths per minute), and normal body temperature (36.5-37.5^0^C) were maintained throughout the experiment through titration of isoflurane dose and with environmental heating.

### Magnetic Resonance Imaging Protocol

In the first set of the experiment (N = 26), 3D-T1w and T1-mapping MR images were taken to visualize CSF-ISF exchange of Gd-chelate into the brain and to quantify Gd-chelate kinetic parameters. The experimental timeline is shown in **Fig. 2A** and Supplementary **Fig. S1, S2**. Detailed imaging protocol is described in Supplementary Methods. For quantitative measurements T1-map RARE protocol was set based on a RARE-sequence with one echo image, RARE factor = two and six T1 experiments. Each experiment has a different TR producing one image. By default, a T1-map is generated automatically for a single slice. Typical values for T1 of the rat brain can be found in previous publications (Behroozi et al., 2018; Guilfoyle et al., 2003; de Graaf et al., 2006). To achieve enough signal to noise ratio within a particular part of the organ, it is recommended to acquire several images so that they cover a time up to five times the T1 (Chow et al., 2012; de Graaf et al., 2006; Guilfoyle et al., 2003; Lin et al., 2000). To ensure agreement of T1-values with the literature, we first optimized the T1-mapping sequences within the hippocampal region of the brain at different spin-lattice relaxation time as shown in **Table S1**. Since T1-values at constant volume of the rat brain were not affected by different spin-lattice relaxation time, we decided to use a TR = 3000ms, 8 min scan time for further studies. A post-processing macro Fitinlsa which was implemented on the 7T Bruker Scanner to automatically start the T1 parameters map calculation to extract quantitative measurements of T1-mapping.

## Analysis

T1 was calculated from the Image Sequence Analysis (ISA) functiont1sat:

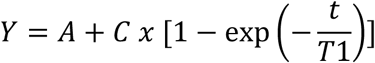

The parameters are defined in the following way:

- A – absolute bias,
- C – signal intensity,
- T1 – spin-lattice relaxation time.

This function supplied by Bruker uses a repetition time list calculated from the protocol parameters to generate the T1 relaxation curve. The fit is based on the magnitude image of the reconstructed dataset. OsiriX 10.0.5 was used to calculate the Gd-enhanced brain volume over time after cisternal Gd-chelate injection using the T1-mapping sequences.

### Focused Ultrasound System

For good acoustic coupling of the FUS beam and for getting access to the cisterna magna for intrathecal injection, the dorsal scalp fur in the sonication trajectory plus up to 3 cm towards neck was removed using standard hair removal cream. After the cisternal injection, animals were placed in a plastic stereotactic frame that is coupled to an MR compatible FUS system (Image Guided Therapy - IGT, Pessac, France), and immobilized with ear bars and a bite bar. A thin layer of ultrasound gel was applied to pair the water-filled coupling membrane of the FUS transducer to the skin of the head. Once the anesthetized animal was secured in the holder and the transducer was placed over its head, approximately at the center of the brain, the assembly was inserted in the bore of the MRI scanner. The sonication trajectory was selected using the remote positioning capabilities of the transducer in all three axes. Stereotactic coordinates for sonication are shown in **Fig. 1B**, *right*. Briefly, an 8 × 10 mm black-dotted rectangle centered on the brain is selected with corners starting from 4 mm lateral at the bregma region (4,0) and move 8 mm to left at (−4,0) then 10 mm posterior at (−4, −10) and then move 8 mm to the right at (4, −10) as indicated in **Fig. 1B**. Ultrasound was held on continuously while the transducer slowly moved around the FUS trajectory for 10 min. Total time to complete one trajectory loop: 24 sec, with a 50 ms pause time between each loop, with 25 total loops for each rat. For sham procedures, the same positioning and trajectory was chosen but the power to the ultrasound transducer was disconnected. FUS (0.25 MI *in situ*, ∼ 7.7% duty cycle for 10 min) or sham was applied transcranially throughout the brain. The orange rectangle represents the expected ultrasound exposure zone based on the ultrasound transducer’s focal spot size (2.78 × 12 mm) that covers a significant part of the rat brain. To account for skull attenuation, a 30% pressure insertion loss was assumed for this size and age of rats (O’Reilly et al., 2011). The expected volume coverage region in rat brain by a single sonication using this particular transducer can be envisioned based on our previous paper **(Supplementary Fig. S1)** (Wang et al., 2018).

### Fluorescence Imaging

In the second set of experiments (N =13), two different sized molecules, free dye (IRDye800CW: 1 kDa) and IR dye-conjugated antibody (Panitumumab-IRDye800: 150 kDa) were co-delivered with Gd-chelate to model the delivery of similar-sized therapeutic agents. The experimental timeline can be found in **Fig. 3A**. Briefly, dyes were diluted with Gd-chelate and injected intrathecally at 0 min. A 3D T1w image was used to confirm the cisternal injection and FUS (0.25 MI *in situ*, 2% duty cycle for 10 min) or sham was applied transracially throughout the brain. About 2 hours after the intrathecal delivery, animals were euthanized with an overdose of euthasol. Then the rat brain was flash frozen in dry ice with 2-methylbutane (Fisher, Pittsburgh, PA). For tissue sectioning, the frozen rat brain was mounted with a minimal amount of optimal cutting medium (OCT) compound and sectioned at a 20 µm thickness using a cryostat (LEICA CM 1950, Buffalo Grove, IL, USA)). Every 10^th^ section (200 μm apart) was saved for optical imaging. The specimen temperature was set at −19 °C and the chamber temperature at −20 °C. Tissue sections were thaw-mounted on microscope glass slides (Fisher, Pittsburgh, PA), fixed the tissue-slides using 4% paraformaldehyde, and applied DAPI for fluorescence imaging. All fluorescence images were collected in a near-infrared fluorescence imager (Ex/Em: 785/820 nm, Pearl Trilogy Imaging System, LI-COR) with 85 µm resolution and processed with Image Studio (version 5.2, LI-COR).

### Hematoxylin and Eosin Staining Histology

In the third set of the experiment (N = 3), T2*w images, as well as hematoxylin and eosin (H & E) staining, were used to evaluate for parenchymal damage. The experimental timeline is shown in **Fig. 4A**. Two hours after the Gd-chelate injection, animals were sacrificed and the brain fixed via transcardial perfusion (0.9% NaCl, 100 mL; 10% buffered formalin phosphate, 250 mL). The brain was then removed, embedded in paraffin, and serially sectioned at 5 μm in the axial plane (perpendicular to the direction of ultrasound beam propagation). Every 50^th^ section (250 μm apart) was stained with H&E.

### Quantification and Statistical Analysis

All MR-images were analyzed in OsiriX (version 10.0.5). An axial plane of T1-mapp sequence with 5/1700 as a lower/upper threshold was used for manual ROI segmentation to calculate the volume of brain parenchymal penetration of the gadolinium tracer. All IRDye images were analyzed in Image Studio (version 5.2, LI-COR). Four-five slices were included from each animal to the analysis. Signals above the background was used for manual ROI segmentation to calculate the area of brain parenchyma penetration of the dye. All the data that were generated from the imaging software were plotted using Microsoft Excel (version 16.16.22). All values were presented as mean ± standard deviation. Statistical analyses were performed with Microsoft Excel (version 16.16.22) and JMP (version 13.2.1). Two-tailed paired Student’s t-test was used to compare the gadolinium-enhanced volume and IRDye-enhanced area between sonicated and non-sonicated (Sham) groups. One-way ANOVA with post-hoc Tukey-Kramer tests were used to compare gadolinium-enhanced volume at different time points (12min, 35 min, 70 min, 105 min, 180 min, 240 min) within the same group, either Ultrasound or Sham. P-values < 0.05 were considered statically significant.

## Supplementary Materials

Fig. S1. Optimization of Magnetic Resonance Imaging sequences.

Fig. S2. Ultrasonic glymphatic induction is safe

Table S1. MRI protocol used for the experiment.

Video V1. 3D visualization of contrast diffusion in MRI.

References

## Acknowledgments

We would like to thank L. de Lecea, K.B. Pauly, L. Fenno, the Stanford Center for Innovations in In vivo Imaging (SCI^3^)-small animal imaging center (NIH S10 Shared Instrumentation Grant S10RR026917-01, PI: M. Moseley), Stanford Animal Histology Services, and the whole Airan lab for helpful discussions and use of equipment. We would also like to thank F. Habte, A. Pascal-Tenorio, L.J. Pisani, T. Doyle, N. Hosseini Nassab for assistance in training and implementation of the varied experimental and image processing protocols. We would like to thank A. Thomas for assistance in figure preparation. This work was funded with an anonymous donor to the Stanford Department of Radiology. Fellowship funding was provided by the Stanford Cancer Imaging Training program (SCIT; NIH T32 CA009695; M.A.).

## Author Contributions

M.A. performed all animal and MRI imaging experiments, ex vivo sample preparation, and imaging data analysis. Q.Z. completed the optical imaging experiments and with E.L.R. provided the IR800-panitumumuab tracer. R.D.A. helped conceive and with M.A. design the overall study, and secured funding for the work. All authors together drafted, edited, and reviewed the figures and manuscript.

