## Supplementary Methods and Figures for "Noninvasive Ultrasonic Glymphatic Induction Enhances Intrathecal Drug Delivery"

### Supplementary Materials

Fig. S1. Optimization of Magnetic Resonance Imaging sequences.

Fig. S2. Ultrasonic glymphatic induction is safe.

Table S1. MRI protocol used for the experiment.

Video V1. 3D visualization of contrast diffusion by MRI.

References.

### Magnetic Resonance Imaging

Magnetic Resonance Imaging (MRI) (an actively-shielded Bruker 7T horizontal bore scanner (Bruker Corp, Billerica MA), with International Electric Co. (IECO) gradient drivers, a 120mm ID shielded gradient insert (600 mT/m, 1000 T/m/s), AVANCE III electronics; 8-channel multi-coil RF and multinuclear capabilities and volume RF coils; and the supporting Paravision 6.0.1 platform) was used to visualize glymphatic transport of Gd-chelate into the brain and to make quantitative T1-maps. The facility provides isoflurane anesthesia in medical-grade oxygen, and physiological monitoring of the subject including ECG, pulse oximetry, respiration, and temperature feedback for core body temperature maintenance by warm airflow over the animal. Following cisterna magna contrast agent injection, an MRI compatible animal FUS system (Image Guided Therapy - IGT, Pessac, France) was used in all experiments. Animals were placed in a prone position in a plastic stereotactic frame that has a single channel radiofrequency transmit-receive head coil (IGT) which is coupled to the FUS system. Animals were immobilized in the frame with ear bars and a bite bar. Noninvasive, MRI-compatible monitors for the respiratory rate and body temperature were used during the imaging session. Following localizer anatomical scout scans, a 3D T1-weighted (T1w) fast low angle shot (FLASH) sequence was acquired in the coronal plane with a repetition time (TR) = 50 ms, echo time (TE) = 3.53 ms, flip angle (FA) = 20°, number of acquisition (NA) = 1, field of view (FOV) = 60 × 40 × 15 mm, slice thickness (ST) = 0.3 mm, total scanning time = 6 min 37 s, acquisition matrix size of 256 × 128 × 128 interpolated to 256 × 256 × 256, yielding an image resolution of 0.313 × 00.267 × 0.313 mm. A standard T1-map rapid acquisition with relaxation enhancement (RARE) with TE = 7ms, TR = 300, 600, 1000, 1500, 2000, or 3000 ms, FOA = 35 x 31 mm, ST = 1mm, scan time = 8 min 58 s, image resolution = 0.273 x 0.242 mm was acquired in a coronal plane to quantify Gd-diffusion within the brain parenchyma. Before setting up that standard T1-map sequence in the study, two more T1-map sequences were taken by replacing the last inversion time, 3000 ms by two different longer times, 4000 ms and 6000 ms to compare how T1-values changes between with cisternal injection of Gd **(Table S1).** Longer sequences did not make a significant change in T1-values at a constant volume so the standard sequence with last TR = 3000 ms was used throughout the study (**Supplementary Fig. S1**). T2* map-multiple gradient echo (MGE) weighted (T2*w) imaging with TR = 796.76ms at different time TE = 3.50, 8.5, 13.5, 18.5, 23.5, 28.5, 33.5, 38.5, 43.5, 48.5 ms, FA = 50°, FOV = 60x40 mm, ST = 1mm, scan time = 4min 44 sec with image resolution = 0.234 x 0.156 mm was used to image whether petechiae, which can result from excessive FUS exposures, occurred **(Fig. 4)**. The scanning protocol consisted of the localizer, baseline, and post-FUS scans of 3D T1-FLASH, T1-map, and T2*w followed by intrathecal administration of Gd-chelate (MultiHance, gadobenate dimeglumine; Bracco Diagnostics Inc, NJ USA; 0.21ml/kg) and/or co-delivery of Gd-chelate with optical tracers **(Table S1)**. A total of 80 µl of the solution was delivered intrathecally using a 27-gauge butterfly catheter (SAI Infusion Technology) within a minute and the first baseline MRI acquisitions were imaged 12 min after the injection. FUS treatment was applied at 23 min after the intrathecal injection. All of the MRI acquisitions continued over either 2, 4, or 72 hours.

**T1-mapping**

As in the Methods, we first optimized the T1-mapping sequences using the values measured of the hippocampal region of the brain with different spin-lattice relaxation times. We found T1-values across a constant volume (0.33 $cm^3$) within the hippocampal region of the rat brain to be 1235 ± 657, 1299 ± 502, and 1298 ± 618 with the longest TR = 3000, 4000, and 6000 ms respectively **(Table 1).** These T1-values are similar to T1-values that are published already in the literature for this particular magnetic strength, 7T (Behroozi et al., 2018; de Graaf et al., 2006; Guilfoyle et al., 2003)**,** and are relatively similar to each other **(Supplementary Figure S1).** To minimize MRI scan time, we decided to use the protocol with 8 min total scan time and TR = 3000ms for the rest of these studies. As observed in the plot in **Figure 2D**, the mean gadolinium-enhanced volume in the sham cohort was 0.17 ± 0.084, 0.25 ± 0.15, 0.22 ±0.1, 0.23 ±0.11, 0.11 ± 0.12 and 0.06 ±0.04 and in the Ultrasound cohort was 0.18 ± 0.07, 0.43 ±0.16, 0.44 ±0.17, 0.45 ± 0.25, 0.18 ± 0.24 and 0.17 ±0.24 at 12 min, 35 min, 70 min, 105 min, 180 min and 240 min respectively.

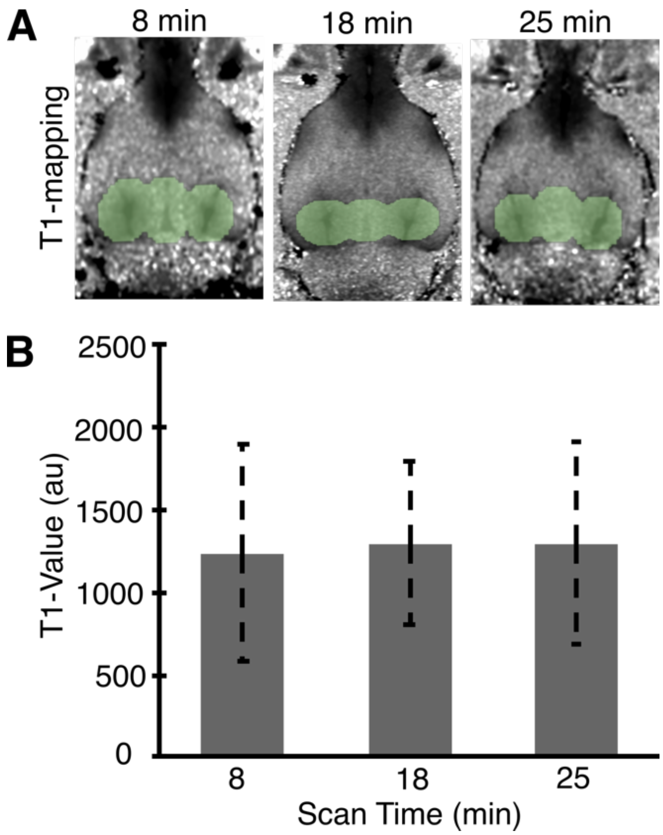


**Fig. S1. Optimization of T1-mapping sequence.** Measured T1 values within the hippocampal region of the rat brain are not affected by TR ranging between 3000 – 6000 ms corresponding to 8-25 min long scan times respectively. **A.** Representative examples of T1-maps with the TR = 3000 ms for 8 min scan time, 4000 ms for 18 min scan time, and 6000 ms for 25 min scan time, each taken at 12 min after intrathecal contrast agent injection. Green circles within the hippocampal region are included for the T1 value comparisons. Constant volumes were used across the different sequences for the T1 value comparisons. **C.** Averaged T1 values across a constant volume (0.37 $cm^3$) for different T1-mapping sequences show that T1 values are not affected by changing TR across these different sequences.

**Ultrasonic glymphatic induction is safe**

The penetration of the MRI tracer into the brain and the presence or lack of petechiae were confirmed using contrast-enhanced T1w and T2*w MRI, respectively **(Figure 4B and S2). Supplementary Figure S2A** shows the parenchymal uptake of Gd-chelate represented by a pseudo-color T1w MR image. The long-term effects of the ultrasound intervention would show on T2*w MR images for up to 72 hours. We observed in the T1w images in **Figure 2B** that the Gd-chelate cleared from the CSF-ISF spaces by 3 hours. Here, we further confirmed that there was no any evidence of Gd deposition **(Figure S-2A)** at 24-72 hours. T2*w revealed that there was no long-term adverse effect such as edema or hemorrhage after the brain-wide ultrasound exposure with 0.25 MI *in situ*, ~7.7% duty cycle for 10 min **(Figure S-2B).**

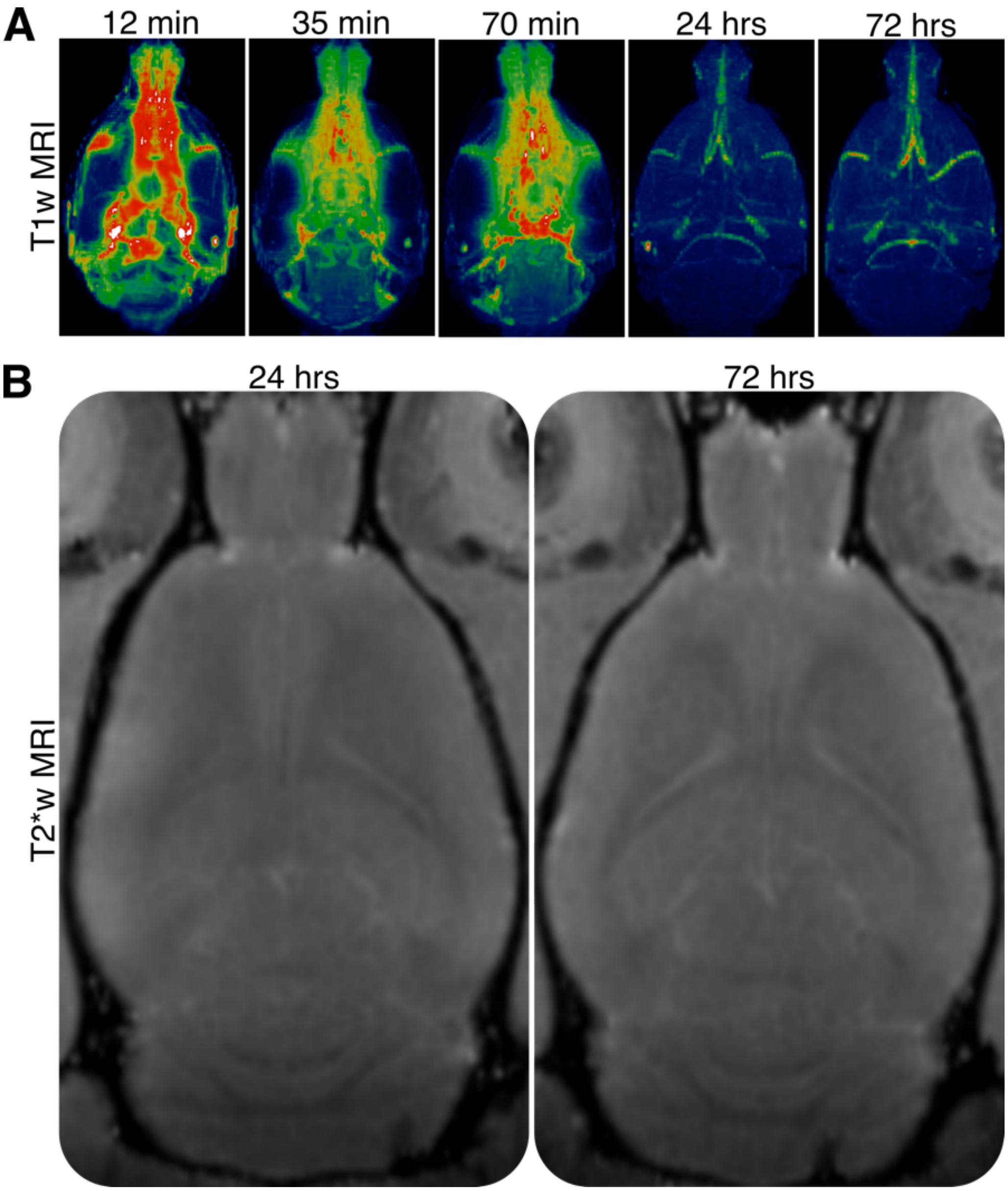


**Fig. S2**. **Ultrasonic glymphatic induction is safe. A.** Representative pseudocolor T1w images of rat brains after cisternal MRI contrast agent injection. Contrast agent uptake was monitored up to 72 hrs given that the CSF is replaced completely by 72 hrs. These results showed that Gd-chelate cleared from the CSF-ISF spaces by 3 hours, and further confirm there is no long-term effects of these interventions. **B.** T2*w MRI showed no brain parenchymal damage up to 72 hours after intervention. No effects such as edema or hemorrhage were noted at 24 hours (**left**) or at 72 hours (**right**) following ultrasound (0.25 MI *in situ*, 7.7% local duty cycle for 10 min) application.

<table>
<tr><th>Name</th><th>Type</th><th>TE (ms)</th><th>TR (ms)</th><th>FA</th><th>FOV (mm)</th><th>ST (mm)</th><th>Scan Time (min)</th><th>Image Resolution (mm)</th></tr>
<tr><td>Localizer</td><td>3 Plane</td><td>2.50</td><td>26.07</td><td>30</td><td>80x80</td><td>1</td><td>0.06</td><td>0.313x0.313</td></tr>
<tr><td>FLASH-T1-3D</td><td>Coronal</td><td>3.53</td><td>50.00</td><td>20</td><td>60x40x15</td><td>0.3</td><td>6.37</td><td>0.313x0.267x0.313</td></tr>
<tr><td>T2*map-MGE</td><td>Coronal</td><td>3.50,8.5,13.5,18.5, 23.5, 28.5, 33.5, 38.5, 43.5,48.5</td><td>796.76</td><td>50</td><td>60x40</td><td>1</td><td>4.44</td><td>0.234x0.156</td></tr>
<tr><td rowspan="3">T1-map RARE</td><td rowspan="3">Coronal</td><td rowspan="3">7.00</td><td>300,600,1000, 1500,2000, 3000</td><td rowspan="3"></td><td rowspan="3">35x31</td><td rowspan="3">1</td><td>8</td><td rowspan="3">0.273x0.242</td></tr>
<tr><td>300,600,1000, 1500,2000, 4000</td><td>18</td></tr>
<tr><td>300,600,1000, 1500,2000, 6000</td><td>25</td></tr>
</table>

**Supplementary Table S1: MRI protocol used for the experiments.** MRI protocol used for each experiment on a Bruker 7T MRI with an IGT single channel transmit/receive coil. *TE: Echo Time; TR: Repetition Time; FA: Flip Angle; FOV: Field of View; ST: Slice Thickness; RARE: Rapid Acquisition with Relaxation Enhancement; FLASH: Fast Low Angle Shot; MGE: Multiple Gradient Echoes*

**Video V1. Caption: 3D MRI visualization of brain contrast agent uptake.**

Representative pseudocolor T1w MRI images of rat brains before and 70 min after sham (top) or ultrasound (bottom) intervention. The hotter colors indicate more MRI contrast agent concentration.
